# FlyBox: A Flexible Open-Source Behavior Monitoring System

**DOI:** 10.1101/2024.05.15.594443

**Authors:** Jasmine Quynh Le, Albert D. Yu, Xihuimin Dai, Christopher Allum, Olivia Jo Bradley, Sophia McAneney, Florian Schwarzinger, Zachary Sherman, Jose E. Alvarez, Gillian Berglund, Daniel Shin, Evelyn Keefer, Lawrence Neeley, Michael Rosbash

**Affiliations:** Howard Hughes Medical Institute and Department of Biology, Brandeis University, Waltham, Massachusetts 02453, USA; Olin College of Engineering, Needham, MA, 02492; Independent Researcher

## Abstract

Over the past two decades, the vast majority of circadian behavior in *Drosophila* has been recorded in Drosophila Activity Monitor (DAM) boards. Though simple and robust, locomotor behavior recording via DAM boards can be prohibitively expensive, especially when taking incubator costs into consideration. Furthermore, their simplicity limits their experimental options and resolution. Here, we present the FlyBox: a simple, open-source benchtop locomotor activity recording system. FlyBox was designed to monitor activity in animals loaded into a standard laboratory multi-well plate. It features light-tight construction and multiple programmable LEDs for simulating day/night cycles and optogenetic manipulation. In total, a single FlyBox costs approximately $750 to build and around two days of labor. In addition, we also present the FlyBoxScanner software to simplify activity monitoring while maintaining compatibility with DAM analysis software. FlyBox is an attractive and affordable package for behavior monitoring that also offers considerable room for customization. Materials and instructions for the FlyBox are available at https://github.com/Rosbash-Lab-FlyBox/FlyBox, and FlyBoxScanner is available at https://github.com/jose-elias-alvarez/flybox-scanner.

## INTRODUCTION

In about 2015, a very talented and original Chinese post-doc Fang Guo in the Rosbash lab came up with the idea for this FlyBox system. The principle is a simple one: it consists of placing single flies in the wells of 96 well plates and monitoring their activity with a video camera. Fang was inspired by the standard video recording of single zebrafish larvae swimming behavior, which he had seen in the literature. He built a similar system for flies and used it for a variety of purposes. They include activity monitoring, the basal requirement, and his box also included lights for entrainment as well as LEDs for optogenetic manipulation; the fly cuticle is so thin that exposing the fly to green or red light is sufficient to manipulate the activity of discrete brain neurons that express channelrhodopsins (Araujo et al., 2020; Guo et al., 2016, 2017).

The Rosbash lab was very keen to develop FlyBox with the broader *Drosophila* community in mind, and we tried twice over several years to develop a production system that could assemble many boxes in a reproducible way. First Fang and a Brandeis undergraduate Hyung Jae Jung, known colloquially as Jae, attempted to build FlyBoxes. (Jae is currently a student at Tufts Dental School.) Fang and Jae as well as our subsequent in-house effort were aided immeasurably by two Brandeis engineers housed in the Physics Department, Greg Widberg and Artie Larsen. They were responsible for transforming Fang and Jae’s initial vision and individual efforts into a functional product.

Jae was very excited about starting a company to sell FlyBoxes, whereas MR was less interested in selling them but keen to see if the system might appeal beyond research laboratories, for example to biology labs for teaching purposes. These labs could be in universities or even in high schools. Doing a simple genetic cross with classic circadian mutants and examining the resultant behavior would be a very simple and rewarding exercise. At one point we brought a FlyBox to Waltham High School to demonstrate neuroscience principles including optogenetics.

However, we were unable to generate anything resembling an assembly line to build multiple FlyBoxes. There were many sources of failure, but one was certainly a lack of sufficient attention on the part of Fang and Jae. Fang was understandably much more focused on his research work, which was essential for him to find a job, and Jae was a rather distracted undergraduate. This description is more critical than it should be, and neglects at least one additional variable (see below); nonetheless, these initial efforts failed beyond the building of a few FlyBoxes. Fang took a couple of them back to China, and I believe he is still using them in his independent laboratory.

We made a second attempt at an assembly line shortly thereafter with a new team and approach. Albert Yu was a fairly new graduate student and teamed up with an advanced undergraduate Max Song to direct a group of undergraduates to build FlyBoxes during the summer. Both Albert and Max were interested more generally in aspects of design and engineering, building things writ large, especially Albert. Albert, like Jae, learned from the engineers Widberg and Larsen. However this effort also failed, for reasons not unrelated to those that sunk our first attempt, namely, Albert and Max were otherwise preoccupied with their principal activities and couldn’t give FlyBox the attention it needed. In addition, we were all deficient in the management skills needed to direct an undergraduate construction assembly line.

More importantly perhaps, the more experience we acquired in building FlyBoxes made it clear that they had multiple design flaws that needed fixing. For example, the boxes were not sufficiently light-tight, and they tended to overheat. The latter was a fan issue. These two problems interfered with one of our major FlyBox goals, namely, a bench-top device that could work without the pricey and cumbersome incubators needed for light and temperature control.

In addition, none of us involved with FlyBox had any idea what it means to design with robust scalable production in mind. Although our prototypes were flawed, they did work but only with a bit of chewing gum and bailing wire. However, scaling is a completely different beast that blindsided us when we began producing multiple FlyBoxes. This issue dwarfed our prototype design flaws. This scaling problem is extremely common for people aspiring to produce an engineered product for market, e.g., see Kickstarter for many examples. One of us (ADY) had been a Kickstarter enthusiast and realized that we had fallen into this same trap.

These failures and other distractions put our FlyBox efforts on hold until one day in late 2021 I began to discuss them with my Brandeis colleague and friend Seth Fraden. Seth is a professor of physics at Brandeis and is in charge of our new engineering program at Brandeis. He listened to my story and suggested I contact the Olin College of Engineering. It is located in Needham, a stone’s throw from Brandeis’ Waltham campus. Seth said that they have a wonderful program called SCOPE, in which a small team of students work together on a project during their entire senior year. So I contacted Olin in the spring of 2022, and we organized a SCOPE effort during the 2022-2023 academic year to design and build a totally new FlyBox.

The SCOPE project required a payment of $50,000 to Olin, my second HHMI payment of exactly $50,000 for activity monitoring. The first one was in 1989, 35 years ago, to TriKinetics for the R&D on the activity monitoring devices that they designed and built in collaboration with us at Brandeis.

The SCOPE team consisted of an Olin professor Lawrence Neeley and 5 wonderful Olin students: Olivia Jo Bradley, Chris Allum, Sophia McAneney, Zachary Sherman, and Florian Schwarzinger. They included all manner of expertise, from mechanical engineering to electronic hardware to software. They worked one day/week on our project throughout the entire academic year, and interfaced every week with three people from my lab: Albert Yu, a post-doc Xihumin Dai and a graduate student Jasmine Le. All 9 of these people are authors of this paper. FlyBox was redesigned from the bottom up, and its current parts list is manageable. All major problems were solved with only minor ones remaining.

During the current academic year, 2023-2024, we worked out some bugs and the lab has now built about a dozen new FlyBoxes. They have been built by 3 different lab individuals: Gillian Berglund, Evelyn Keefer and Daniel Shin. They are also coauthors. Our current estimate is that it takes between 1 and 2 days for one person to build a single FlyBox; it costs about $750 for parts. Although FlyBox still has some modest problems, it works well as a benchtop locomotor activity monitoring device.

In addition to the physical FlyBox, we present a new Python-based software to simplify motion tracking and data acquisition. Previous iterations of the FlyBox integrated multiple software packages to perform image acquisition, well segmentation, and conversion into a DAM-compatible output format for downstream analysis. Our new FlyBoxScanner software package integrates these disparate steps into one, by live-populating an output file during an experiment; this file is compatible with existing downstream analyses.

Together, we present an open source suite of hardware and software to facilitate, reduce the cost and democratize activity monitoring in circadian biology.

## RESULTS

### FlyBox experiments are an accessible method for quantifying activity-based behaviors

The fundamental concept of the FlyBox is simple. Fruit flies - or other organisms of interest - are loaded into a standard laboratory multi-well plate, which is then placed into an enclosed chamber equipped with an infrared camera to record locomotor activity. The camera is connected to a computer and records on command via a software - discussed below - which tracks single organisms within individual wells (Figure 1). The use of multi-well plates allows for flexibility in number of fruit flies observed, size of arena, and accommodating larger organisms that may require more space such as mosquitos. These plates have been standardized in scientific research for many years for cross-platform compatibility and are readily available from many scientific manufacturers. For many longer timescale experiments using flies, it is standard practice to provide food and water for longevity of the organisms. Each well of a plate can be filled with an agar gel with water and sucrose; we find that 2% agar and 5% sucrose in water works well and most flies survive for more than two weeks. Using an agar gel also allows for clear recording of behavior with the infrared camera in the FlyBox. We have validated the use of FlyBox for locomotor activity-based behavior, which can be extrapolated to sleep behavior as well.

**Figure 1.**
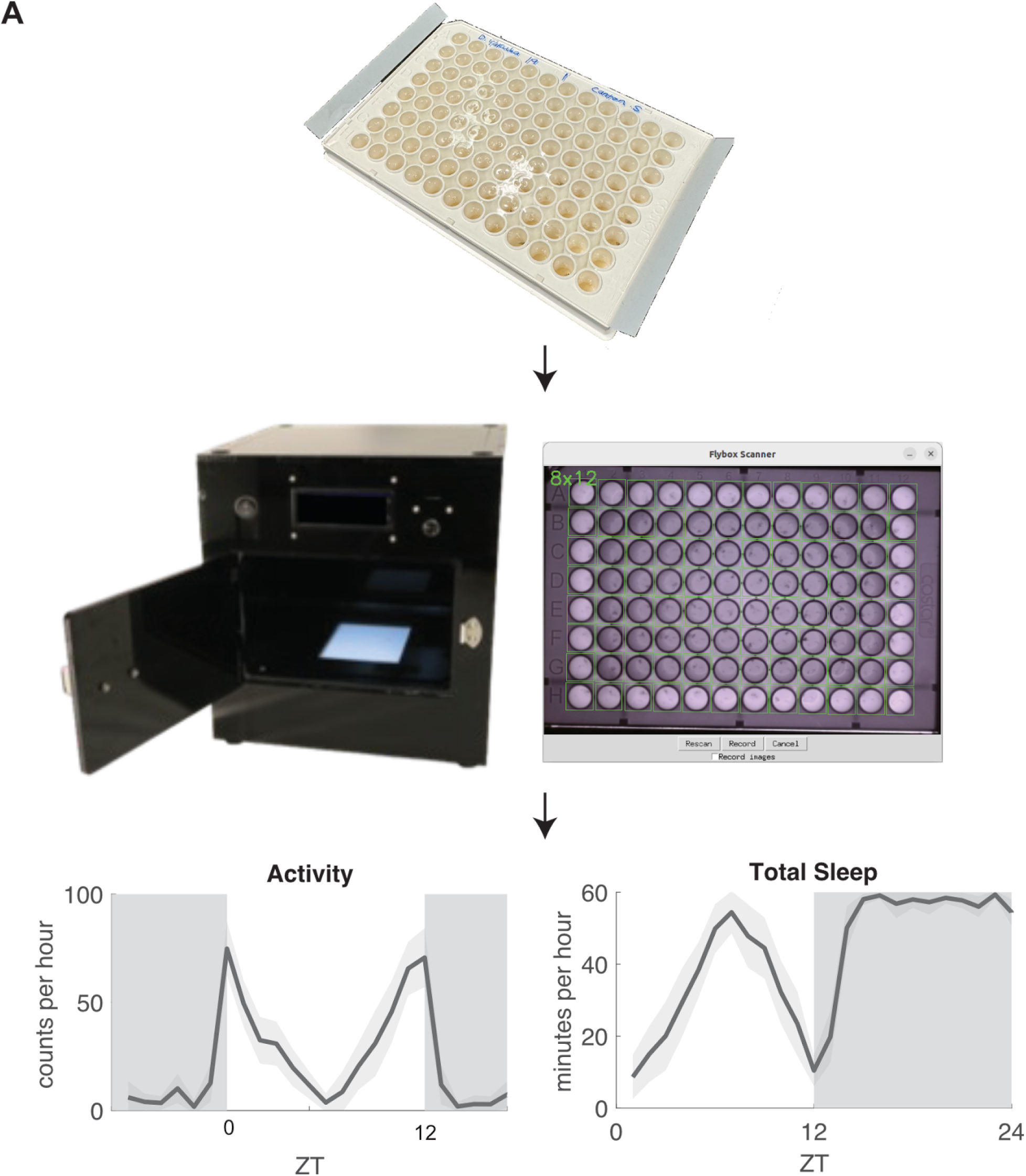
FlyBox experiments are an accessible method for quantifying activity-based behaviors. A) (Top) Standard laboratory plates are loaded with food and flies. (Middle) Plates with animals are loaded into FlyBox, which is connected to a computer with FlyBox Scanner software. (Bottom) Typical output data is locomotor activity and sleep.

### FlyBox is an open-source, low-cost, and easy build

Our open-source design of FlyBox is meant to be accessible to a wide range of scientific minds: from students in classrooms to researchers at big institutions. FlyBox is a free-standing box with three major compartments (Figure 2A). The top compartment houses an infrared camera that can record animals in light and dark, and an Arduino. It also contains a set of white LEDs that serve as entrainment lights to mimic day and night. Additionally, a set of red and green LEDs can be installed in the top compartment for conducting optogenetic experiments. The top compartment is separated from the middle compartment by a white translucent diffuser panel that allows for even illumination of the middle compartment. The middle compartment features a removable platform that fits a standard multi-well plate, and can be customized at the experimenter’s discretion. Below this platform is the bottom compartment, which contains a set of infrared LEDs that provide constant infrared illumination throughout the duration of an experiment.

**Figure 2.**
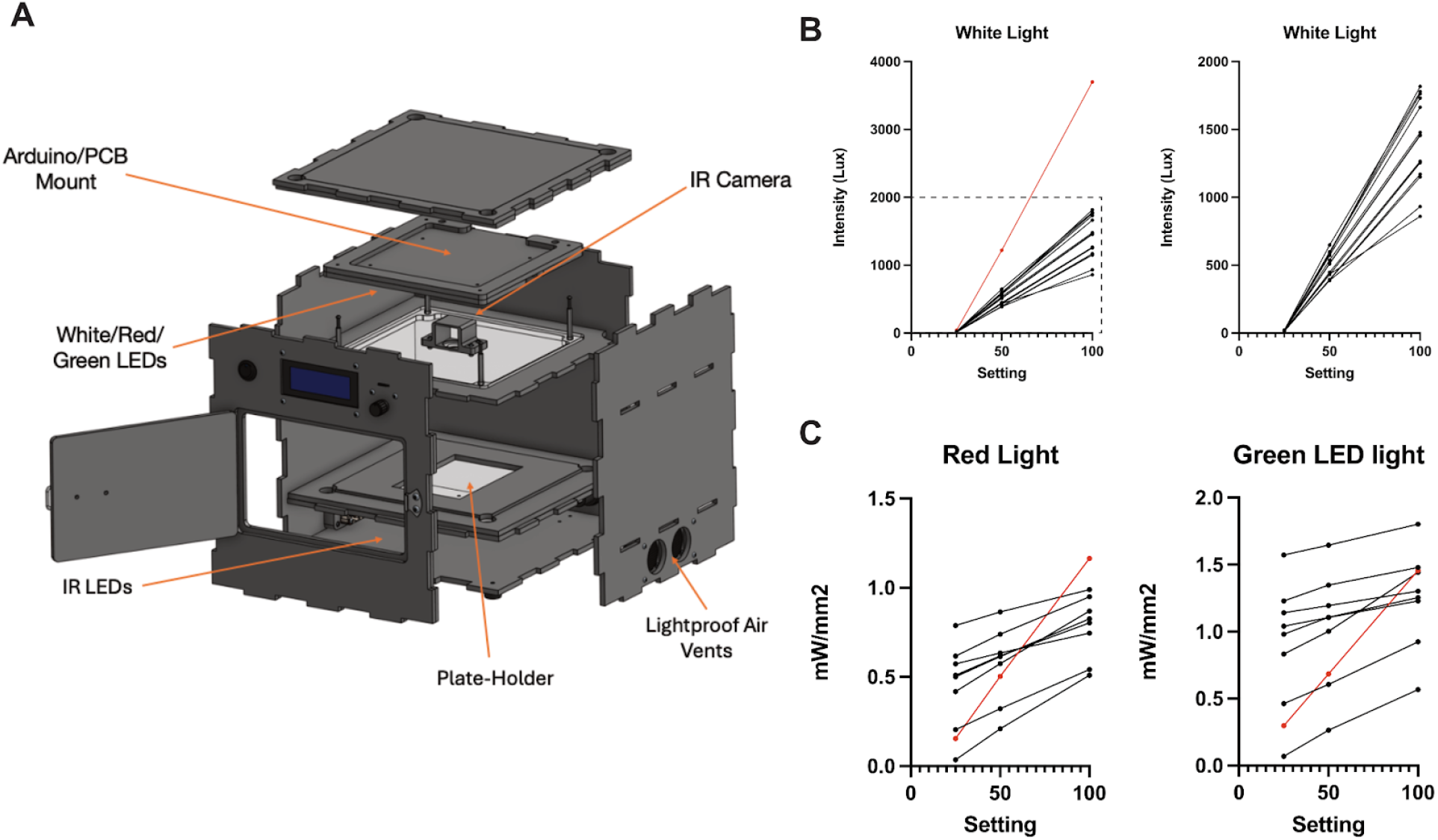
FlyBox is an open-source, low-cost, and easy build. A) FlyBox consists of three main compartments. The top compartment houses the Arduino and LED lights. The middle compartment is where the plates are loaded and consists of the stage where plates are laid. The bottom compartment houses the IR lights, which allow for recording of animals in darkness. B) Light intensity curves for the white LEDs in lux. (Left) Black lines represent intensity values for FlyBoxes with a white diffuser panel and the red line represents the intensity values for a FlyBox with a clear diffuser panel. (Right) Inset of dashed box on the left. Intensity values for FlyBoxes with a white diffuser panel. C) Light intensity curves for the red (left) and green (right) LEDs in mW/mm2. Black lines represent intensity values for FlyBoxes with a white diffuser panel and the red line represents the intensity values for a FlyBox with a clear diffuser panel.

The intensity of the IR LEDs are controlled by a rotary encoder, which is set after building to the user’s preference and is otherwise largely untouched. The white, red, and green LEDs are controlled by the Arduino through custom firmware. We provide a simple web application (https://rosbash-lab-flybox.github.io/FlyBox/) that enables a user to control the timing, intensity, and frequency of the white, red, and green LEDs. Despite our best efforts for consistent builds, we have found that there is variability between the intensity output of each FlyBox with the same intensity setting. With the white LED, we found that with intensity setting 25, the output range is 13-21 Lux, while intensity setting 100 has an output range of 860-1800 Lux with our standard white diffuser panel (Figure 2B, right). We also measured one FlyBox with a clear diffuser panel instead of a white diffuser panel; the intensity output values were about twice as high at setting 50 and 100 with the clear diffuser panel (Figure 2B, left). For the red LED, setting 25 measured at 0.035-0.8 mW/mm2 while setting 100 measured at 0.5-1 mW/mm2, and for the green LED, setting 25 measured at 0.07-1.6 mW/mm2 while setting 100 measured at 0.6-1.8 mW/mm2. With a clear diffuser panel, the red and green LED were approximately within the same range as the white diffuser panel, however the incline with increased intensity setting was much steeper. All FlyBoxes include previously reported ranges for entrainment light intensity and ontogenetic light intensity for CsChrimson (Klapoetke et al., 2014) and GtACR1 (Mauss et al., 2017). Given these results, it is important to consider inter-FlyBox variability when using multiple FlyBoxes for an experiment. Additionally, if higher white light intensity is required for an experiment, a clear diffuser could be used, which does not seem to be the case for the red and green LEDs.

### FlyBoxScanner is a new software that detects wells, tracks animals, and records activity to an output file

In addition, we also present a new software package - FlyBoxScanner - to significantly simplify data acquisition and accelerate workflows (Fig. 3A). Previous work conducted in the FlyBox used a combination of periodic image capture and PySolo software to convert time-course images into distance traveled in a DAM-compatible format. FlyBoxScanner simplifies this workflow considerably.

**Figure 3.**
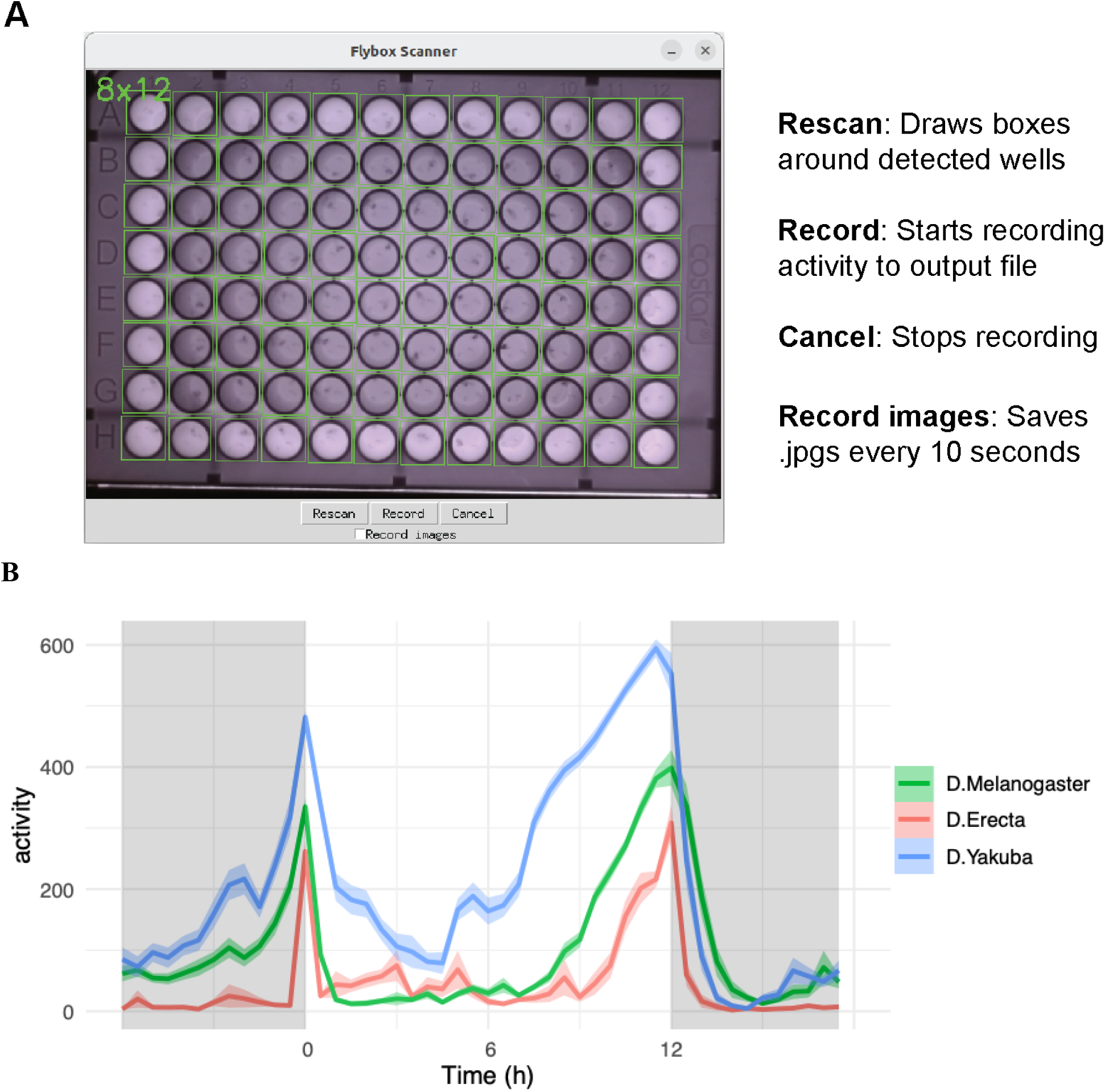
FlyBoxScanner is a new software that detects wells, tracks animals, and records activity to an output file. A) Screenshot of FlyBoxScanner software and the functions of available buttons. B) Behavior of 3 different Drosopholids over 3 LD days, averaged, recorded using FlyBoxScanner software.

The FlyBoxScanner software uses the OpenCV package to automatically segment wells and track behavior. First, it uses a circle detection algorithm to segment the frame into a grid of wells. Next, it performs background subtraction to detect the flies as contours, then assigns each fly to a well within the grid. As new frames are generated, each frame is compared to the previous frame where a contour was detected in the same well. The delta between contours is then reported as motion in the output file. It is identical to the output file of DAM boards, except that it has as many columns as there are wells instead of the fixed 32 from TriKinetics. FlyBoxScanner also features tunable sensitivity to suit an individual user and/or applications needs.

To test FlyboxScanner, we examined the LD behavior of three different Drosophilids - *D. melanogaster, D. yakuba*, and *D. erecta* (Fig. 3B). FlyBoxScanner faithfully recapitulates bimodal activity exhibited by all three Drosopholids.

### FlyBox can be used to study circadian activity and sleep patterns using white entrainment lights

*Drosophila* are crepuscular animals, exhibiting bimodally dispersed activity in the early morning and in the late evening. This behavior is preserved when subject to constant darkness (Renn et al., 1999), while any semblance of rhythmicity is largely abolished in constant light (Emery et al., 2000). FlyBox was designed to be light tight, so that experiments can be done with the entrainment lights serving as the Zeitgeber on the bench top. When we entrain wildtype w1118 flies in a FlyBox with a 12 hour light-dark cycle (12:12 LD), we see the expected bimodal activity pattern with higher levels of activity around ZT00 and ZT12 (Figure 4A, top panel labeled LD, light blue). We also observe expected sleep patterns, which includes higher levels of sleep midday during the siesta and even higher levels of sleep throughout the night (Figure 4A, bottom, LD, light blue). Both of these patterns are dampened but persist in constant darkness (Figure 4A, DD1-DD4, light blue). It is evident that there is an activity peak that remains close to ZT12. With the classic per short mutants that are known to have short periods, we also observe a shorter free running period in constant darkness, as shown by the activity peaks that start earlier with each subsequent DD day (Figure 4A, red). Also consistent with decades old literature, we observe that wild-type flies that are entrained to 12:12 LD in FlyBox (Figure 4B, LD) slowly lose rhythmicity in first days of constant light and become arrhythmic by day 4 of constant light (Figure 4, LL). This is true for the two wild-type strains we tested, w1118 and CantonS. A third method of utilizing white light is for sleep deprivation. Like humans, flies sleep mostly throughout the night when it is dark. When we entrain wild-type CantonS flies in 12:12LD and then have one day of LL, we find that flies are more active and sleep less between ZT12 and ZT18 compared to flies that remain in 12:12LD (Figure 4C). These experiments show that FlyBox recapitulates known circadian behavior recorded in DAM boards.

**Figure 4.**
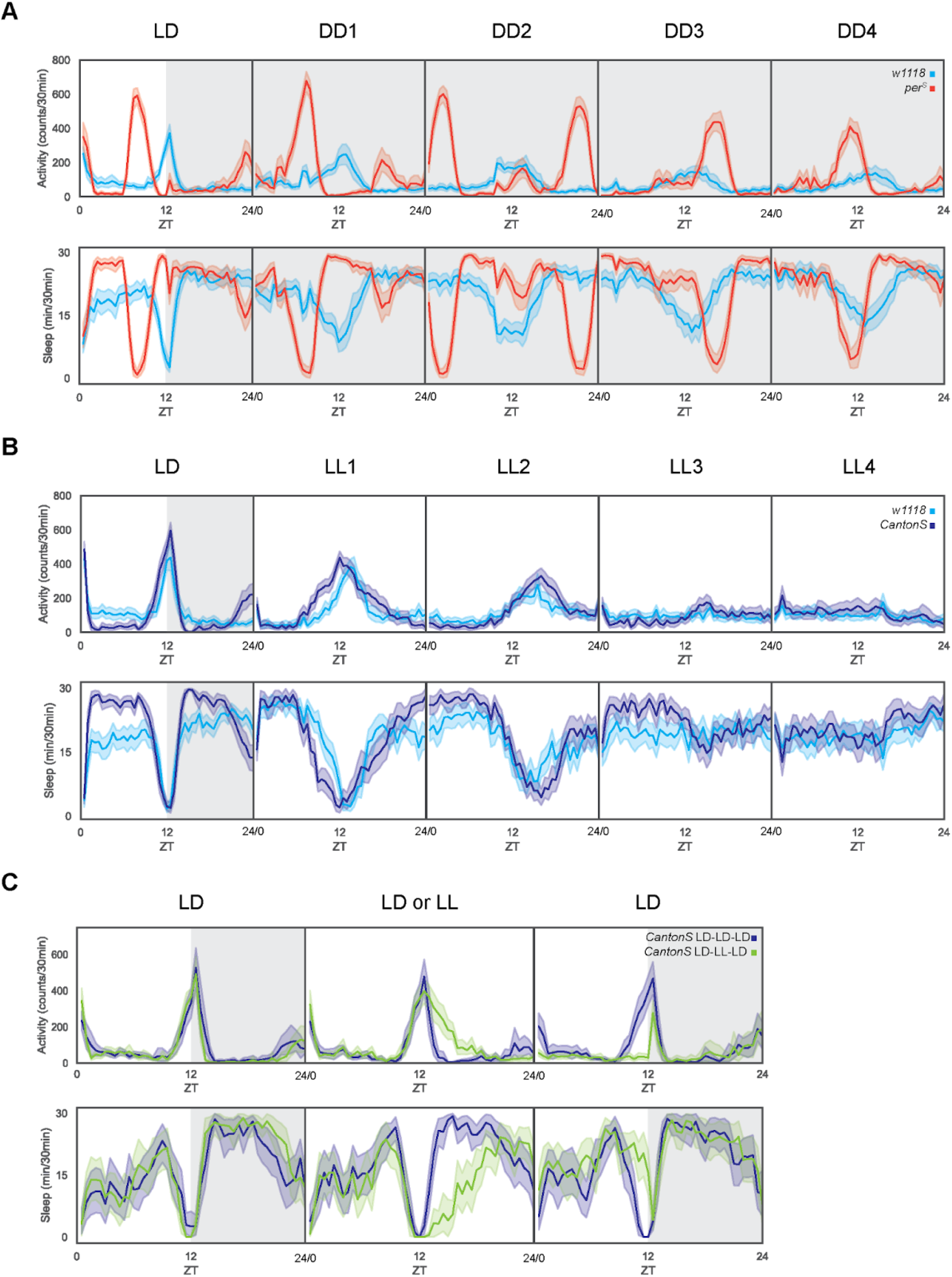
FlyBox can be used to study circadian activity and sleep patterns using white entrainment lights. A) Example constant darkness experiment with wild-type w1118 flies. Flies are entrained in 12:12 LD and then put into constant darkness (DD) for four days. B) Example constant light experiment with wild-type CantonS (blue) and w1118 (orange) flies. Flies are entrained in 12:12 LD and then put into constant light (LL) for four days. C) Example sleep deprivation via light experiment with wild-type flies. Flies were either exposed to 12:12 LD for three days (gray) or exposed to 12:12 LD on day 1, constant light (LL) on day 2, and 12:12 LD on day 3 (blue).

### FlyBox is equipped with red and green LEDs for optogenetic manipulation

The FlyBox is also equipped with red and green LEDs for the purpose of optogenetic neuronal excitation and inhibition. We tested two light-sensitive ion channels - CsChrimson, which is sensitive to red light and drives depolarization upon red light exposure, and GtACR1, which are chloride channels sensitive to green light that silence neurons upon exposure. To test the effectiveness of red light stimulation in FlyBox, we had experimental flies that express UAS-CsChrimson under a tissue-specific promoter or control flies that had UAS-CsChrimson and an empty GAL4. Both strains were entrained to 12:12LD and exposed to red light for 24 hours starting at ZT00. With red LED illumination, the control flies exhibited more activity and less sleep in comparison to the day before due to the red light keeping flies awake in the dark (Figure 5A, gray). The experimental flies, however, had a much smaller evening activity peak and higher levels of sleep during red LED illumination, meaning neuronal excitation, compared to the day before (Figure 5A, pink). This indicates that excitatory activity of these neurons promotes sleep. We also tested the effectiveness of green light optogenetic inhibition using experimental flies expressing UAS-GtACR1 under the control of a different tissue-specific GAL4 and control flies that had heterozygous expression of either the UAS or the GAL4. All three strains were entrained to 12:12LD before being exposed to green light for 24 hours starting at ZT00. Similar to the effect of red light on control strains, green LED illumination caused more activity and less sleep on control strains in comparison to the day before due to light exposure during dark (Figure 5B, gray). In the experimental flies, green light illumination caused a decrease in evening activity and an increase in nighttime sleep in comparison to the day before (Figure 5B, green). This suggests that these neurons are typically activity promoting when not inhibited. Both excitatory and inhibitory experiments here illustrate that the red and green LEDs in FlyBox can be used for optogenetic manipulation.

**Figure 5.**
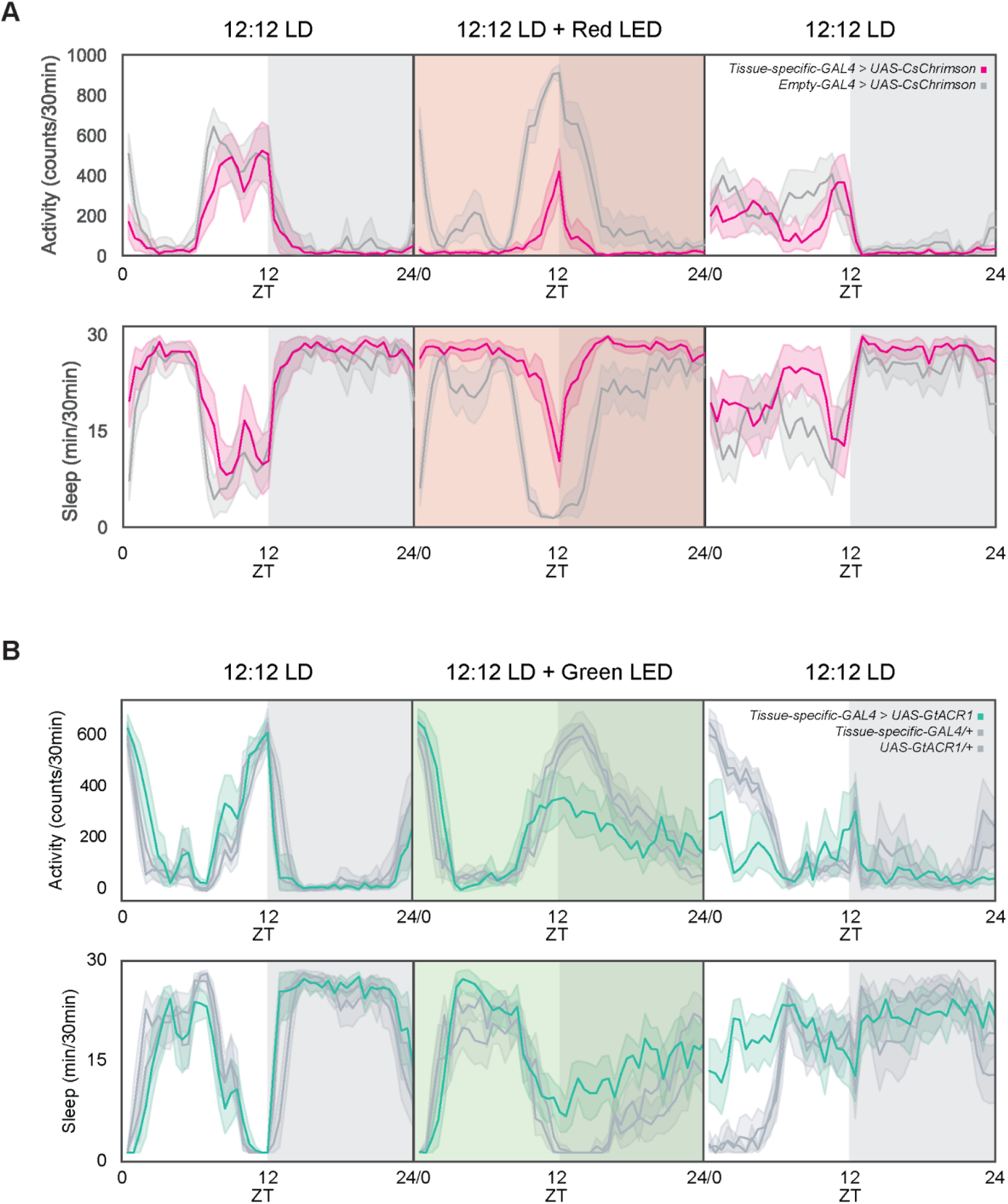
FlyBox is equipped with red and green LEDs for optogenetic manipulation. A) Example optogenetic activation experiment with red light-sensitive channelrhodopsin CsChrimson. Flies with neurons labeled by a GAL4 driver and expressing UAS-CsChrimson are in red and heterozygous parental controls are in gray. Top row of data is activity in counts per minute and bottom row is sleep data in minutes of sleep per 30 minutes. Left panel is the baseline day with 12:12 LD, the middle panel is 12:12 LD with red light illumination, and the right panel is 12:12 LD. B) Example optogenetic inhibition experiment with green light-sensitive channelrhodopsin GtACR1. Flies with neurons labeled by a GAL4 driver and expressing UAS-GtACR1 are in green and heterozygous parental controls are in gray. Top row of data is activity in counts per minute and bottom row is sleep data in minutes of sleep per 30 minutes. Left panels are the baseline day with 12:12 LD, the middle panel is 12:12LD with green light illumination, and the right panels are 12:12 LD with green light illumination.

## DISCUSSION

Here, we present FlyBox: a behavior monitoring system made open to the scientific community. The open-source nature of the FlyBox opens it up to user-specific preferences and applications. We have already experimented with many modifications - including swapping LED colors, modifying the FlyBox to support two plates instead of one (for 192 flies), using anywhere between 6- to 96-well plates for different sized insects, and potentially also applications requiring higher resolution behavior analysis. We believe that flexibility presented by the FlyBox will enable many new avenues of experimental investigation.

## DATA AVAILABILITY

Materials and instructions for the FlyBox are available at https://github.com/Rosbash-Lab-FlyBox/FlyBox, and FlyBoxScanner is available at https://github.com/jose-elias-alvarez/flybox-scanner.

The most up to date bill of materials is available at: https://docs.google.com/spreadsheets/d/1IHYl5jYujjvnLfjMplQ0Q5b2qjvNmh8gb_7PU2YNHrg/edit#gid=1894107250

And the most up to date assembly instructions are available at:

https://docs.google.com/document/d/1mAbPdf_ScTb2pJxWNEVFJWNXVA8fg-mKAZQye8iWYWE/edit

